# Patterns and Predictors of Tic Suppressibility in Youth with Tic Disorders

**DOI:** 10.1101/204933

**Authors:** Christine A. Conelea, Brianna Wellen BA, Douglas W. Woods, Deanna J. Greene, Kevin J. Black, Matthew Specht, Michael B. Himle, Hanjoo Lee, Matthew Capriotti

## Abstract

Background: Tic suppression is the primary target of tic disorder treatment, but factors that influence voluntary tic inhibition are not well understood. Several studies using the Tic Suppression Task have demonstrated significant inter-individual variability in tic suppressibility but have individually been underpowered to address correlates of tic suppression. The present study explored patterns and clinical correlates of tic suppression in youth with tic disorders using a large, pooled dataset.

Methods: Individual-level data from 9 studies using the Tic Suppression Task were pooled, yielding a sample of 99 youth with tic disorders. Analyses examined patterns of tic suppressibility and the relationship between tic suppressibility and demographic and clinical characteristics.

Results: A large majority of youth demonstrated a high degree of tic suppression, but heterogeneous patterns of tic suppressibility were also observed. Better tic suppressibility was related to older age and more frequent tics but unrelated to other clinical variables, including presence of psychiatric comorbidity, psychotropic medication status, and tic and premonitory urge severity.

Conclusions: The mechanisms underlying the observed heterogeneity in tic suppressibility warrant further investigation. The Tic Suppression Task is a promising method for testing mechanistic hypotheses related to tic suppression.

---

Tic disorders (TDs) are childhood-onset conditions characterized by motor and/or phonic tics [1]. Unlike symptoms of many other movement disorders, tics associated with TDs are experienced as “unvoluntary” [2] in that they can be inhibited via prefrontally-mediated cognitive control mechanisms [3]. The vast majority of patients describe tics as occurring in response to premonitory urges, aversive somatosensory experiences that are temporarily alleviated by tic execution [4,5]. Tic suppression is the primary target of behavioral, pharmacological, and neurostimulation treatments [6,7]. Successful voluntary tic suppression underlies the effectiveness of empirically-supported behavioral interventions for TDs, such as Comprehensive Behavioral Intervention for Tics (CBIT; [6]), which teach specific skills for inhibiting tics in the presence of premonitory urges.

Given the central importance of voluntary tic inhibition to nonpharmacological tic management, researchers have studied various questions related to tic suppression using the Tic Suppression Task (TST) developed by Woods and Himle [8] (also see [4,9–15]). In this paradigm, a participant is seated in front of a computer or token dispenser and told it is a “tic detector.” In reality, an experimenter unobtrusively observes the child (e.g., using a one-way mirror), and tic frequency is measured via direct observation coding. This deception is employed to reduce reactivity effects, as observation can have significant and non-uniform effects on tic expression [16].

Standard TST conditions include baseline (BL; instruction to let tics occur freely) and reinforced suppression (SUP; instruction to suppress tics plus reward delivered by “tic detector” after every 10 sec tic-free interval). Additional conditions have been added to this basic paradigm to test experiment-specific hypotheses. Typically, participants are exposed to brief segments of each condition (e.g., 5-10 min) in alternating or random order, with multiple iterations of each condition. The key dependent variable is tic frequency, calculated as a mean rate of tics per minute for each condition. This design allows for within-subject analyses of tic rate by condition and group-level analyses to describe aggregate patterns across conditions. The first study using the TST (n = 4; [8]) demonstrated a substantial decrease in tic frequency from BL to SUP (76.3% reduction), which far exceeded the ~10% decrease from BL to “verbal instruction” condition (instructions to suppress without supporting reward contingency). Subsequent research has repeatedly and reliably replicated this effect of reward-enhanced suppression on tic frequency (e.g., [17–19].

Despite robust findings of reward-enhanced suppression at the group level, significant inter-individual variability in degree of tic suppressibility has been noted. Reasons for this heterogeneity are unknown. Individual studies using the TST have been underpowered to assess correlates of suppression because of reliance on small samples (N*s*= 4-15), reflecting feasibility limitations inherent in studying a low-prevalence disorder. Researchers have theorized, or indirectly demonstrated, that tic suppressibility heterogeneity may be related to demographic and clinical factors, such as age, premonitory urge severity, and co-occurring psychopathology [18,20].

Age and premonitory urge severity have been theorized to impact tic suppression, but supporting evidence is limited. Some have suggested that older children may be better able to suppress tics due to increased capacity to sustain attention to effortful tasks [3,8]. Consistent with this, Banachewski, Woerner, and Rothenberger [21] found that older youth were more likely to self-report an ability to suppress tics (age 15-19 years > 11-14 years > 8-10 years). However, the age-tic suppression relationship has yet to be tested using objective tic frequency data. Premonitory urges may facilitate tic suppression by serving as a cue to engage in suppression [5]. Experimental research on the urge-tic temporal relationship has confirmed that urge intensity increases prior to tics and that tics are more likely to occur when urges are higher [22,23]. Whether urges are a prerequisite for successful suppression is unclear, with one study in adults demonstrating no association between video-rated tic suppression capacity and premonitory urge severity [24].

Somewhat more evidence from studies using the TST suggests a possible relationship between tic suppression and attention problems (e.g., those related to attention deficit hyperactivity disorder; ADHD), although findings are mixed and derived from small samples. Significant associations have been found between decreased tic suppressibility and increased attention problems on a parent-rated scale [25] and continuous performance task [18]. Interestingly, Greene et al. [12] reported significant correlations between tic suppression and parent-rated attention problems but in the opposite direction, such that those with more inattention symptoms demonstrated greater tic suppression.

Although individual studies using the TST have advanced our understanding of the factors that influence tic suppression, clinically relevant questions about patterns and predictors of suppression have been difficult to address within single, small studies. The present study explored patterns and clinical correlates of tic suppressibility in youth with TDs using a large, pooled data set comprised of data from multiple laboratory studies using the TST. First, we sought to describe the group-level effect of suppression contingent reward on tic frequency and the distribution of tic suppressibility across individuals. Second, we aimed to evaluate candidate demographic and clinical correlates of tic suppressibility. Based on prior research, we hypothesized that better tic suppression would be positively associated with older age and premonitory urge intensity, while poorer tic suppression would be associated with an ADHD diagnosis.

## Methods

*Study selection*. A literature search in databases PubMed and PsycINFO identified research studies utilizing the TST [8] that included conditions of BL and SUP. Fourteen studies were identified and authors were contacted to request individual level, de-identified data for published studies or other protocols their lab had disseminated via peer-reviewed abstracts/conference presentations. All authors responded to our inquiry. Data were available for 9 studies [4,9–12,14,18,26,27]. Individual-level datasets were no longer available for older studies [8,13,17,20,25]. A summary of included studies is presented in Table 1. The current study was determined by the Lifespan-Rhode Island Hospital Institutional Review Board (IRB) to not qualify as human subjects research due to the use of de-identified data only.

*Participants*. To be included in current analyses, individual participants must have met DSM-IV-TR or DSM-5 criteria for a tic disorder (Tourette Disorder, Chronic/Persistent Tic Disorder, or Transient/Provisional Tic Disorder) and had available a Yale Global Tic Severity Scale (YGTSS; [28]) total severity score and tic frequency scores (tics per minute) for conditions of BL and SUP. A total of 99 youth met inclusion criteria.

*Measures* were selected based upon common availability across studies. Demographic variables included age, gender, and medication status. Medication status was available for n = 97. Due to study variability in medication tracking methods, medication status was coded as presence vs. absence of 1) any psychotropic medication, 2)antipsychotic, 3)stimulant, 4)α_2_ adrenergic agonist, or 5) serotonin reuptake inhibitor (SRI).

Psychiatric diagnostic status was determined using well-validated, structured clinical interviews (see Table 1). Psychiatric comorbidity data were available for 87 youth (excluding [18]) and grouped into the following major diagnostic categories: 1) anxiety disorders (generalized anxiety, separation anxiety, social anxiety, specific phobia, panic, and post-traumatic stress disorders), 2) obsessive-compulsive disorder (OCD), and 3) ADHD (any subtype). Mood disorders (major depressive and bipolar disorders, dysthymia) were not analyzed as only n = 1 had a mood diagnosis. Note that some studies had exclusion criteria related to specific psychiatric diagnoses (see Table 1).

Tic severity was measured using the YGTSS, a clinician-rated assessment of tic severity over the past week that produces an overall tic severity score ranging from 0-50, such that higher scores indicate greater severity [28]. Premonitory urge intensity was measured using the Premonitory Urge for Tics Scale (PUTS; [29]), a self-report measure yielding a total score ranging from 9-26, such that higher scores indicate more intense urge experiences. PUTS scores were available for 88 youth.

Tic frequency scores represent the mean number of tics per minute for all within-subject replications of a given condition (BL or SUP), such that each participant had one mean tic frequency score for each condition. To assess the degree to which youth were able to suppress tics during SUP, we calculated a percent change variable that we refer to here as “tic suppressibility” [(BL tic frequency - SUP tic frequency)/BL tic frequency*100]. Positive values indicate tic reduction during SUP (i.e., better suppression), near-zero values indicate little to no difference between conditions (i.e., poor suppression), and negative values indicate tic increase during SUP (i.e., tic worsening during suppression conditions).

*Data analysis plan*. All measures were examined descriptively. The relationship between tic suppressibility and demographic/clinical characteristics was assessed using independent samples t-tests for dichotomous variables (gender, diagnostic status, medication status).

Nonparametric tests were used when data did not meet t-test assumptions. A two-tailed Wilcoxon signed-rank test was used to compare tic frequencies in BL and SUP. The Mann-Whitney test was used to examine tic suppressibility within specific medication classes given unequal sample size distribution. Pearson’s correlations were used to examine the relationship between tic suppressibility and continuous variables (age, total YGTSS and PUTS scores, baseline tic frequency).

## Results

*Participant characteristics*. Participants had a mean age of 11.2 years (SD = 3.0, range = 5.0 - 17.7). Gender distribution reflected the known preponderance of tics in males (male: n = 78, female: n = 21). Forty-two youth (42.4%) had a co-occurring psychiatric diagnosis. Thirty-three had ADHD, 28 had an anxiety disorder, and 15 had OCD, and 1 had a mood disorder; 21 had more than one comorbid diagnosis. About one-third were taking a psychotropic medication (n = 29, 29.3%); 17 were taking more than one medication. In terms of specific medication classes, 12 (12.1%) were taking an α_2_ adrenergic agonist, 10 (10.1%) an SRI, 9 (9.1%) a stimulant, and 8 (8.1%) an antipsychotic.

Tic severity on the YGTSS fell in the moderate range (m = 22.3, SD = 7.7, range = 7-39). The mean PUTS total score reflected moderate urge intensity (m = 18.1, SD = 6.2, range = 9-31).

*Tic suppressibility*. BL mean tic frequency was 7.6 tics per min (SD = 5.9, median = 6.2) and SUP was 2.6 tics per min (SD = 2.9, median = 0.8). Data were not normally distributed (BL: *z*_skewness_ = 1.5, *z*_kurtosis_ = 2.96; SUP: *z*_skewness_ = 2.1, *z*_kurtosis_ = 4.5; all *p* <.001). Tic frequency was significantly lower during SUP than BL, *T* = 20.7, *z* = -7.9, *p* <.001, and the observed effect size was large, *r* = −.56.

Median tic suppressibility was 71.1% (SD = 38.3), and 89.8% of participants (n = 89) had scores above zero, indicating lower tic rate during SUP than BL. Wide variability was observed (−61.5% through 98.8%; see Figure 1). About 70% (n = 71) of participants had at least a 50% decrease in tic frequency. Notably, 20.2% (n = 20) showed near complete suppression during SUP (90% or greater reduction), 10.1% (n = 10) showed no suppression (tic suppressibility between −25% and 25%), and 7.1% (n = 7) showed tic worsening when trying to suppress (26% or greater increase).

**Figure 1.**
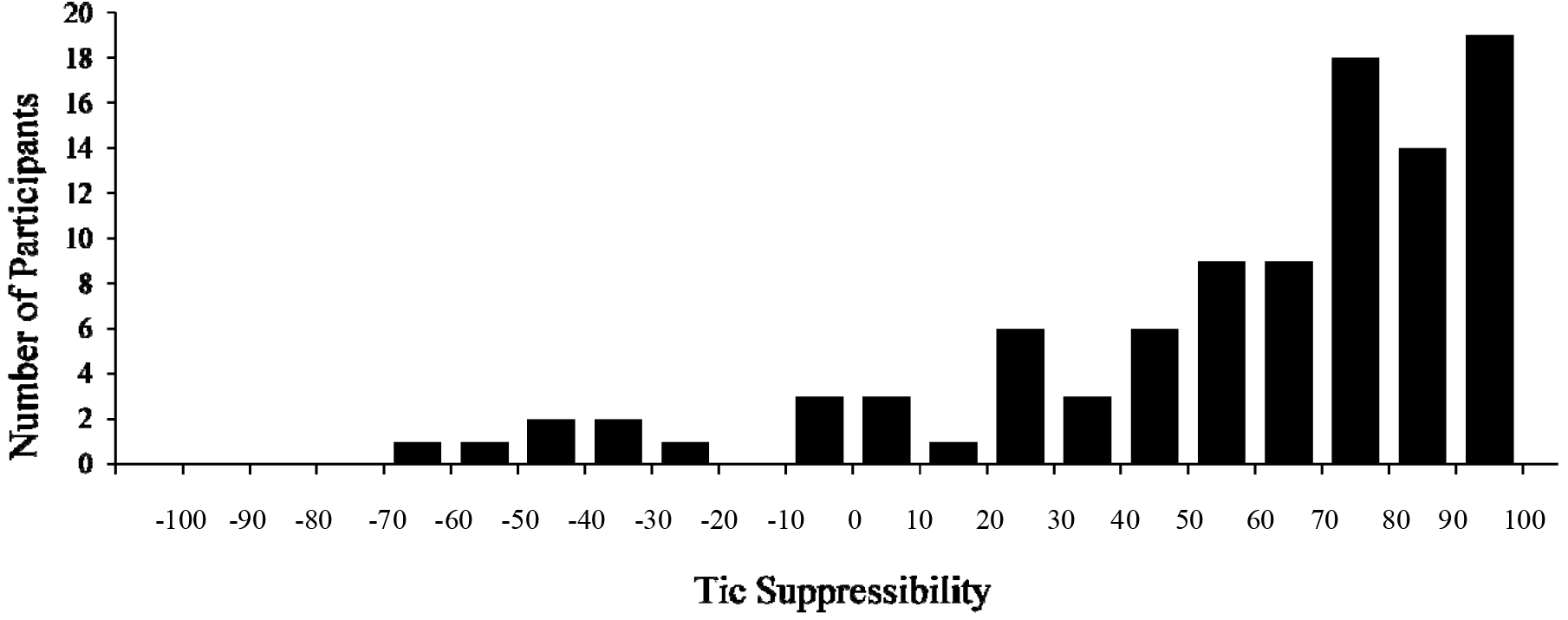
Tic Suppressibility Across Participants.

**Figure 2.**
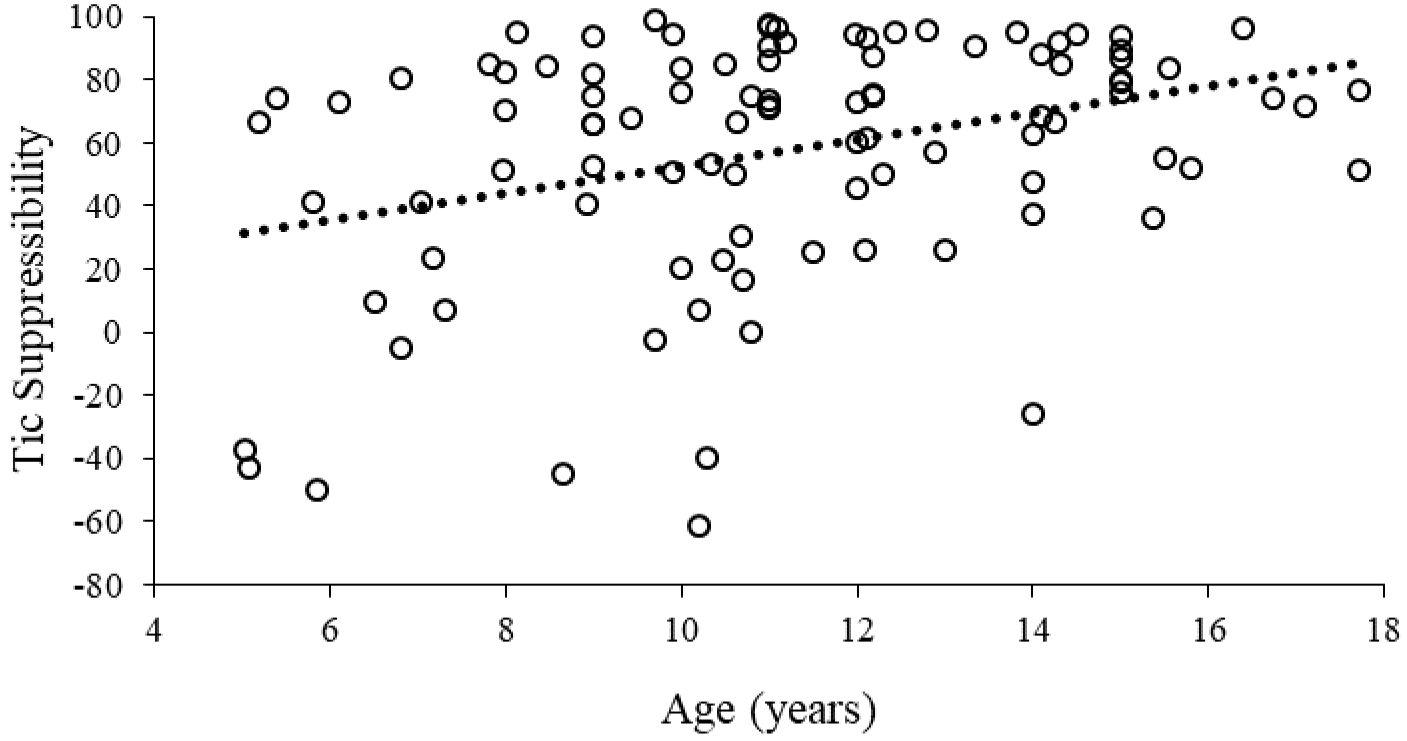
Tic suppressibility as a function of age. Hashed line represents a least-squares regression line fitted to the data.

*Relationship of tic suppressibility to demographic and clinical characteristics*. Age was positively correlated with tic suppressibility, *r* = .34, *p* = .001, such that older children demonstrated a greater degree of tic suppression. There was no gender difference in tic suppressibility, *t* = − 1.19, *p* = .23. Tic suppressibility did not differ based on the presence of any psychiatric comorbidity (*t* = -.28, *p* = .77) nor specifically in terms of ADHD (*t* = .95, *p* = .34), anxiety disorder (*t* = −.02, *p* = .98), or OCD status (*t* = −.43, *p* = .67).

There was no difference in tic suppressibility for those taking vs. not taking a psychotropic medication, *t* = −.73, *p* = .46. No differences in tic suppressibility were found for specific medication classes: antipsychotic (n = 8), *U* = 329.0, *p* = .72; stimulant (n = 9), *U* = 277.0, *p* = .13; α_2_ adrenergic agonist (n = 12), *U* = 431.0, *p* = .38; SRI (n = 10), *U* = 320.0, *p* = .17.

Baseline tic frequencies and tic suppressibility were significantly correlated (*r* = .26, *p* = .007), such that those with more frequent tics at baseline showed greater tic suppressibility. Correlations of tic suppressibility with YGTSS-measured tic severity (*r* = .18, *p* = .07) and with PUTS-rated premonitory urge severity (*r* = .15, *p* = .17) were in the same direction but not significant.

## Discussion

We evaluated reinforced tic suppression and its correlates using data from 99 youth with TDs pooled across 9 studies. Results showed that a large majority of youth achieved robust tic suppression when attempting to suppress tics in the presence of a supporting reward contingency, but significant inter-individual heterogeneity in tic suppressibility was also observed. Results regarding correlates of tic suppression were only partially consistent with a priori predictions: tic suppressibility was positively related to child age (older children tended to suppress more effectively) but not significantly related to premonitory urge severity or ADHD diagnosis.

That robust tic suppression was observed in a majority of participants may be seen as unsurprising, given positive results of prior studies whose data contributed to this study. However, the nature of interindividual variability in tic suppressibility has been difficult to characterize in previous small studies. Though a large majority demonstrated a high degree of tic suppression (i.e., 70% of youth had at least a 50% decrease in tic frequency), we also observed sizable portions of the sample to show particularly striking patterns of suppression, including 20% who had near complete suppression, 10% who seemed unable to suppress, and 7% whose tics *worsened* during SUP.

It is critical to note that results describe tic suppressibility in an austere lab setting with an immediate reward contingency supporting suppression. Thus, the ability of these participants to suppress tics under naturalistic conditions is unknown. Behavioral interventions (e.g., CBIT) focus on tic suppression practice in everyday life; thus, the present findings do not suggest that most youth with tics already have the same tic suppressing repertoire which behavior therapies aspire to develop. Instead, we suggest that most youth naive to behavior therapy can indeed voluntarily suppress tics under certain, “ideal” conditions. Therapy focuses on enhancing suppression abilities and generalizing them to more naturalistic contexts, so that skills may be utilized concurrently with engagement in normal activities and in the absence of contrived reinforcement contingencies. From an experimental perspective, the notion that tic suppression can be experimentally induced for most children confirms the utility of this paradigm for systematic tests of how extraneous factors perturb tic suppression, such as contextual events or medication inductions.

The finding that tic suppressibility improves with age is a novel finding consistent with a neurodevelopmental view of TDs. The ability to suppress interfering actions develops throughout childhood and adolescence and is driven by functional and structural maturation of the prefrontal-striatal network [30]. Although TDs are associated with atypical function in control-related regions [31,32], based on maturation alone we would still expect older youth to have greater top-down control of tics compared to their younger counterparts. The notion that older people with tics develop altered or compensatory neurofunctional organization of motor control networks as a result of tic suppression experience has also been proposed to explain findings that those with TDs tend to show normative or enhanced behavioral performance on motor control tasks [33–35]. Given that suppression occurred in the context of reward, it is also possible that the age finding reflects the particular heightened responsiveness to reward in adolescence vs. both childhood and adulthood, which has been attributed to imbalanced maturation of limbic regions relative to cortical control regions [36]. Alternatively, age-related differences may simply be due to the fact that older children have experienced longer illness duration, lending greater opportunity to naturally develop and practice short-term tic suppression strategies. Testing youth longitudinally with the TST may inform developmental models of tic suppression.

The presence of comorbid psychiatric disorders was not associated with differential tic suppressibility. Although comorbidity has been shown to adversely impact psychosocial functioning in those with TDs (e.g., [37]), the literature has been inconclusive in terms of comorbidity effects specifically on tic suppressibility. Interestingly, although anxiety disorders have been found to predict poorer response to CBIT [38], we found no evidence of impaired tic suppression in youth with comorbid anxiety.

ADHD has been the disorder most frequently hypothesized to be detrimental to tic suppression, based on the rationale that tic suppression involves attentional and inhibitory control mechanisms that are impaired in those with ADHD. Our results are consistent with previous studies showing that youth with ADHD can achieve robust tic suppression [13] and benefit comparably from CBIT [38]. As ADHD is known to have a heterogeneous neurocognitive profile [39], tic suppression may be related to neurocognitive and/or learning processes that are *implicated in* ADHD, rather than related directly to the ADHD clinical phenotype. One interesting consideration is whether suppression contingent reinforcement might mitigate the impact of ADHD on tic suppression, which would be consistent with research showing enhanced sensitivity to reward in children with ADHD [40]. This possibility could be tested by comparing tic suppressibility with and without a supporting reward contingency in children both with and without ADHD. Empirical evaluation of this possibility might guide treatment refinement, as it could indicated whether reinforcement to enhance suppression bolsters treatment response in youth with tics and ADHD.

We also found no difference in tic suppressibility related to psychotropic medication status. This parallels findings from the CBIT trials, which showed that medication status did not moderate treatment response[38], and previous research showing that dexmethylphenidate does not impair tic suppression in youth with tics and ADHD [13]. We did not find evidence of impaired tic suppression among youth taking antipsychotic medication, which is incongruent with findings showing blunted response to CBIT among individuals on these medications [38]. However, null findings in terms of medication class effects should be interpreted with caution given the small numbers of children taking specific medications. It is also notable that medicated youth in this sample still met study entry criteria involving a moderate degree of tic severity, meaning that they could be undertreated or medication non-/sub-optimal responders. Future research could use the TST to more rigorously test whether specific medications impact tic suppression, either in an experimental fashion [13] or as a pre-post assessment when medication withdrawal is not feasible.

We did not find a significant relationship between tic suppressibility and tic severity, as measured by the YGTSS. This finding may reflect the notion that tic generation and tic suppression are separate processes subserved by different loops of cortico-striatal-thalamo-cortical (CSTC) circuits [41]. Findings may also reflect measurement variance. The YGTSS total score is a global rating of multiple dimensions of tic symptomatology over the previous week. Of these dimensions, tic frequency has been the sole dependent variable in TST studies. How suppression efforts impact other symptom dimensions, such as tic intensity or complexity, remains unknown and should be tested in future research. It is notable that a significant correlation was found between tic frequency at baseline and tic suppressibility, such that youth with more frequent tics during BL showed a greater degree of suppression. While it is possible that this finding is a measurement artifact (i.e., more frequent tics yield more opportunity to have a larger change score), it may also be the case that more frequent tics afford more “real life” practice in suppression, which could enhance skills related to suppression (e.g., greater awareness of tic sensations, naturally formed competing motor behaviors) and development of compensatory neurocircuits.

Tic suppressibility was not significantly correlated with PUTS-rated premonitory urge severity. Behavioral models have conceptualized premonitory urges as a facilitator of tic suppression, as they provide salient cues to initiate tic inhibition [6]. In recent years, findings, including those in the current study, call this idea into question [12,22,24,42], instead suggesting that the urge-tic relationship is heterogeneous across individuals and more nuanced than once thought [4,19,23]. For example, in the CBIT trials [38], individuals with higher PUTS scores demonstrated blunted treatment response, raising the possibility that higher urges are associated with a more treatment-refractory form of TD. Neuroimaging studies indicate that distinct neural circuits subserve tic and urge generation [41,43], suggesting that the two processes are at the least dissoci*able*, even if not necessarily dissociated under all conditions.

Our findings have implications for future research. Mechanistic work is needed to identify the reasons for heterogeneous patterns of tic suppression across individuals. Based on our current findings and the prior literature, it now seems clear that variables capturing broad clinical characteristics are not useful as predictors or moderators of tic suppressibility and are therefore unlikely to shed light into the mechanisms underlying suppression or to have prognostic utility. Candidate mechanisms should be explored using fine-grained, transdiagnostic variables that provide better insights into brain-context-behavior interactions that drive tic suppressibility. The TST continues to hold strong promise as a method for probing mechanisms underlying tic suppression, particularly because it is a well-established laboratory task that can be paired with examination of other units of analysis (e.g., neurocircuitry).

Limitations of the current study include those inherent to combining a dataset across studies, such as statistical weaknesses associated with secondary analyses, method variance across projects, possible uncontrolled or unmeasured site effects, and access to only a limited range of variables that were common across studies. We did not have access to information about medication indications, and only a small number of participants were taking medications in each examined medication class. Finally, continuous measures of comorbidity were not available. Although comorbidity rates parallel those reported for clinical trials with similar inclusion/exclusion criteria [44], rates are lower than those expected in the general TD population.

In conclusion, our results indicate that a large majority of youth with tic disorders are able to achieve robust tic suppression in a laboratory tic suppression paradigm. Better tic suppressibility was related to older age and more frequent tics but unrelated to other clinical variables. The mechanisms underlying the observed heterogeneity in tic suppressibility warrant further investigation, as enhancement of tic suppression is a key treatment target. The TST holds promise as a method for subtyping TD samples to more precisely identify mechanisms driving differential tic suppression ability, predicting response to behavioral interventions, or matching individuals to treatment components, consistent with a personalized medicine approach.

## Author’s Roles

All authors were involved in data collection. CC and MC led project conception, organization, and execution; designed the analytic approach; and wrote the first draft. CC conducted data analyses. BW created the study database. All authors edited, reviewed, and approved the final version of the manuscript.

## Financial Disclosures of All Authors for Preceding 12 Months

CC reports grant support from the National Institute of Mental Health. DW reports grant support in partnership with PsycTech for development of www.tichelper.com; grant support and speaker fees from the Tourette Association of America; and royalties from Oxford University Press, Guilford Press, and Springer Press. DG reports grant support from the National Institute of Mental Health. MH reports grant support from the National Institutes of Health and grant support, speaker fees, and honoraria from the Tourette Association of America. MC reports clinical research training funds from the American Academy of Neurology and Tourette Association of America. KB reports grant support from the National Institute of Mental Health, Michael J. Fox Foundation, and Pershing Square Foundation, clinical trials support from Neurocrine Biosciences, Inc., and Sunovion Pharmaceuticals, Inc., and speakers bureau and advisory board compensation from Acadia Pharmaceuticals, Inc. BW, MS, HL and have no reportable disclosures.

